# Atrioventricular Nodal Function, Atrial Fibrillation And Ventricular Tachycardia in Giraffes

**DOI:** 10.1101/2021.01.14.426686

**Authors:** Tejas E. Shivkumar, Joseph Hadaya, Mads Frost Bertelsen, Christian Aalkjær, Morten H. Smerup, Tobias Wang, Olujimi A. Ajijola, Barbara N. Horowitz

## Abstract

Unique cardiovascular adaptations in giraffes (*Giraffa Camelopardalis reticulata*) have been the focus of numerous investigations for almost a century. The vertical distance between the heart and brain impose high pressure on the giraffe left ventricle leading to thickening that exceeds heart in other mammals. Yet, cardiovascular function appears to be unimpacted by these morphologic differences. Physiologically adapted atrioventricular conduction may contribute to these unique cardiovascular characteristics. Atrioventricular (AV) function was assessed to determine whether physiologically adapted AV intervals might optimize the delay between atrial and ventricular contractions. Using ambulatory and intracardiac recordings, we report a longer PR interval in giraffes than predicted by allometric scaling. Slow ventricular response during atrial fibrillation further supports species-specific atrioventricular adaptations.

## Introduction

Unique cardiovascular characteristics of giraffe are associated with the species’ notable left ventricular hypertrophy. The adverse functional consequences of left ventricular hypertrophy in humans may not be present in giraffe (Smerup et al., 2016; Aakljaer and Wang, 2021; Ostergaard et al., 2013). Several adaptations could account for this, including an accommodation for ventricular filling by a physiologically adapted atrioventricular interval. The atrioventricular (AV) conduction time in sinus rhythm, inferred from the PR interval of the electrocardiogram, and intra-cardiac electrophysiological (EP) study provides an objective measurement of the physiological delay between atrial and ventricular contractions. In mammals, an allometric scaling of PR interval as ¼ power of body mass (BM) has been proposed (Noujaim et al., 2004). To explore this further, we utilized a unique opportunity to study three giraffes; using an insertable loop recorder in one giraffe, while invasive electrophysiological measurements were performed in the other two giraffes. We report here, for the first time, ambulatory ECG data and invasive electrophysiological measurements in giraffes. The giraffe has a PR interval in this study longer than the interval predicted by the universal allometric relationships and could reflect its adaptation to its specific physiological adaptation. Induced atrial fibrillation in a giraffe was also associated with a very slow ventricular response.

## Methods and Results

These studies were part of the UCLA-Aarhus joint program on neural control of mammalian hearts.

### Experimental animals

Experiments were performed on three juvenile male giraffes (*Giraffa camelopardalis Linnaeus 1758*, age 25, 30 and 32 months) weighing 465, 510 and 550kg. The experimental protocol was approved by the national Danish Animal Experiments Inspectorate (Danish Ministry of Justice). Local ethical committee members oversaw the experiments and permission to euthanize the animals was granted by Aarhus University.

### Animal handling and anesthesia

All studies were performed on anaesthetized giraffes using an experimental protocol and a set-up as described previously by Brøndum et al. (Brondum et al., 2009); following overnight fast and 2h without water, giraffes were pre-medicated using 50 mg xylazine and guided to a chute where they were blindfolded. Anesthesia was induced by remote injection of etorphine (9 µg kg−1, i.m.) and ketamine (0.9 mg kg−1, i.m.), which rendered the giraffes recumbent within minutes. As soon as the animals were recumbent, a cuffed endotracheal tube (internal diameter, 20 mm) was inserted to allow ancillary ventilation with oxygen using a demand valve (Hudson RCI). After a supplementary dose of ketamine (0.2 mg kg−1, i.v.), the giraffes were placed in right lateral recumbency. Animals were allowed to breathe spontaneously, but were mechanically ventilated, if necessary, to maintain normal end-tidal CO2 and arterial blood gases. Anesthesia was maintained by continuous infusion of α-chloralose (KVL Pharmacy, Frederiksberg, Denmark; 30 mg kg−1 h−1 gradually decreasing guided by clinical signs) into the saphenous vein. Following systemic administration of heparin (150 units kg−1; 25,000 IE ml−1; B. Braun, Melsungen, Germany) and local infiltration by lidocaine (2%; SAD, Copenhagen, Denmark), vascular sheaths were placed in the carotid artery and jugular vein at the base of the neck. The measurements completed in the present study would be virtually impossible to achieve in conscious animals (or even conscious sedation equivalent as done in humans), the necessary restraint and handling stress would be ethically unacceptable and cause considerable disturbance to electrophysiological variables. Anaesthetics also influence the cardiovascular system; however, as mentioned above, the effects of etorphine and medetomidine are considered to be small and α-chloralose is thought to have smaller effects on circulation than most other anaesthetics and has been used by us for a dynamic regulation of anesthesia depth (Covert et al., 1992). Experiments were conducted under experimental animal license (Aarhus University).

**ECG:** needles in skin and placement of electrodes similar to human torso.

### Surgically Implanted Cardiac Monitor

*<u>Giraffe</u> <u>1:</u>* A Medtronic (Reveal LINQ™) Insertable Cardiac Monitor with a wireless transmitter (courtesy Medtronic Inc, Minnesota, MN, USA) was used for ambulatory monitoring.

### Invasive Electrophysiological study

*<u>Giraffe</u> <u>2:</u>* This procedure was done indoors in normothermic conditions. Intracardiac catheters were placed via the left carotid and internal jugular veins. A Millar pressure transducer was inserted via the left carotid artery and placed in LV for pressure measurement.

*<u>Giraffe</u> <u>3:</u>* This procedure was initially planned to be indoors but rain resulted in outdoor procedure, leading to hypothermia (temperature approximately 32-33 degrees. Intracardiac catheters were placed via the left carotid and internal jugular veins. A Millar pressure transducer was inserted via the left carotid artery and placed in LV for pressure measurement. All recordings were recorded on an EP Tracer system (Schwarzer-Cariotek®, Germany) and digital data was stored and reviewed after the study.

### Subject Characteristics and Anesthesia Protocol

Giraffes were pre-medicated using 50 mg xylazine, 50mg ketamine and 40mg scopolamine IM (delivered by remote injection). After 10-20 minutes animals were physically restrained in a restraint device for blindfolding and placement of a head collar for head control during induction. Anesthesia was induced using 3.6mg etorphine and 280mg ketamine IM. Giraffes were endotracheally intubated and breathed 100% oxygen delivered through a demand valve. In giraffe 1, anesthesia was reversed using 7mg atipamezole, 100mg naltrexone and 100mg doxapram IV. In Giraffes 2 and 3, anesthesia was maintained through continued repeated dosing with 0.1mg etorphine and 50 mg ketamine approximately every 20 minutes based on clinical signs.

#### Giraffe 1: Heart Rhythm in conscious giraffe

A loop recorder was implanted in left chest wall subcutaneously (Medtronic Linq™), such that giraffe 1 could be monitored for seven days while it was in familiar surroundings and could interact with other giraffes. The lower and upper cut off for HR were programmed to 60 and 115 bpm, respectively. Sensitivity was programmed to 35 µV with 150 ms blanking after sense and 150 ms sensing threshold decay delay. Average heart rates were 59.0±11.6 and 56.0±8.4 bpm (mean ± SD, range 46-80 and 44-66) and night and day, respectively, daily activity was 98±113 min (mean ± SD, range 23-340 min) and the heart rate variability was 152.0±41.7 (range 89-194).

#### Giraffes 2 & 3: Invasive electrophysiological studies during anesthesia

*<u>Giraffe</u> <u>2</u>*: This procedure was done indoors in normothermic conditions (Figure 2). Two intracardiac catheters with four platinum recording electrodes (Abbott ®) were placed via the left internal jugular veins. Underlying rhythm was sinus. RR interval of 1700 ms, PR interval of 380 ms, QRS duration of 134 ms, QT interval of 612 ms (QTc 469 ms). There were periods of isorhythmic AV dissociation with concealment into the atrioventricular junction. The animal developed atrial fibrillation with catheter manipulation, however, there was a very slow ventricular response RR interval 2086±39 ms (mean±SD). Atrial fibrillation terminated after lidocaine infusion. A His bundle signal could not be recorded with blind catheter manipulation. AV nodal ERP was 1100 ms (with a basic drive train of 1700 ms). AV Wenckebach cycle length was 1400 ms. Ventricular pacing showed no VA conduction at baseline. Following norepinephrine bolus (4 mg single dose) the left ventricular systolic pressure (LVSP) increased to 372 mm Hg from a baseline of 151 mm Hg, and the animal developed AV block with PP interval of 1117±17 ms (mean±SD) and RR interval of 1963±53 ms (mean±SD). The T waves changed morphology from being broad based to being narrower. There was no VA conduction during ventricular pacing after norepinephrine (NE) bolus. Ventricular ectopic beats were seen after NE bolus and sustained ventricular tachycardia was also seen (50 seconds) that spontaneously terminated.

**Figure 1:**
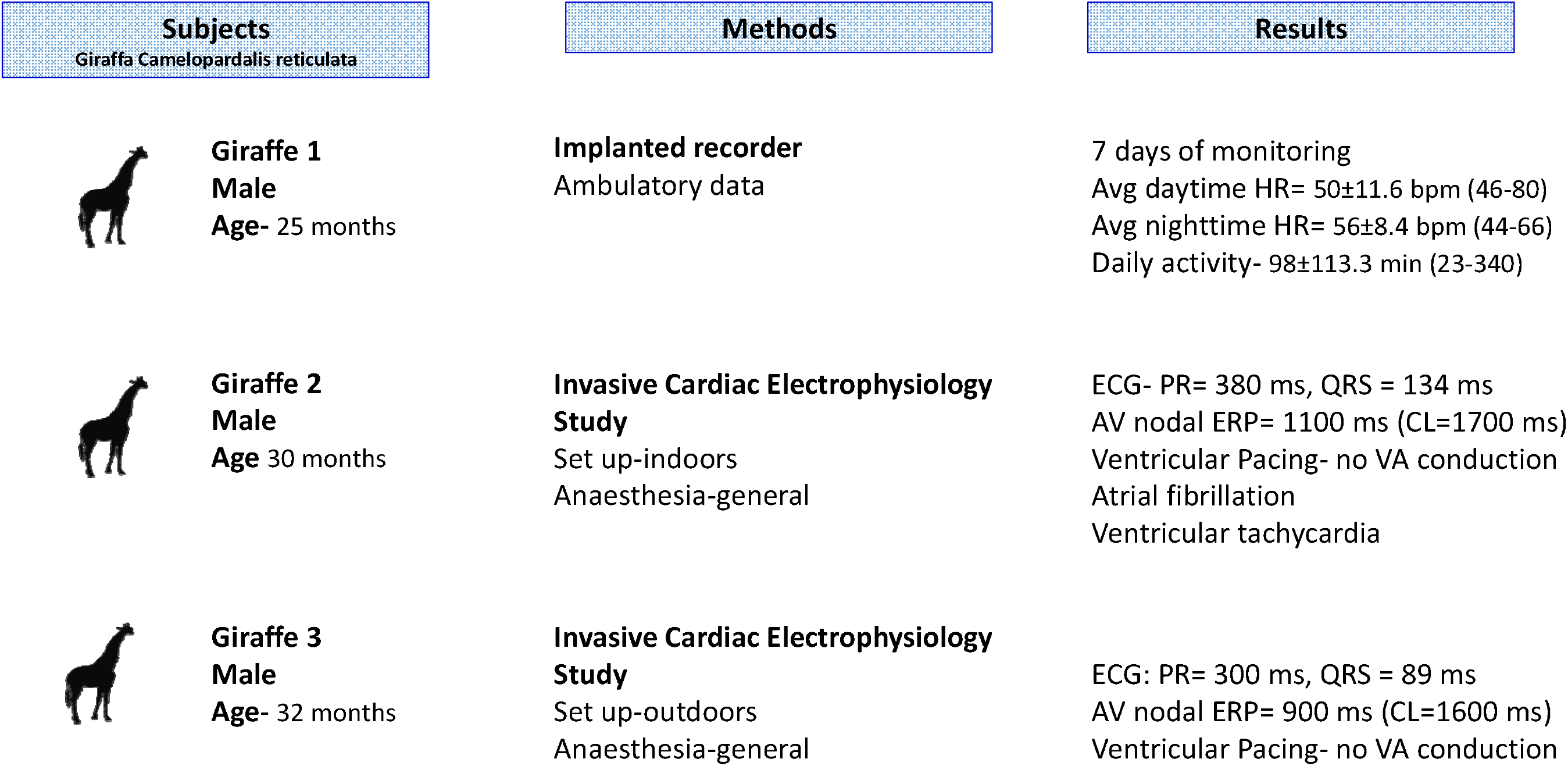
Methods & Results. ECG= electrocardiogram, AV= atrioventricular, HR= heart rate, ERP= effective refractory period, CL= cycle length, ms= millisecond, bpm= beats per minute, VA= ventriculoatrial

**Figure 2:**
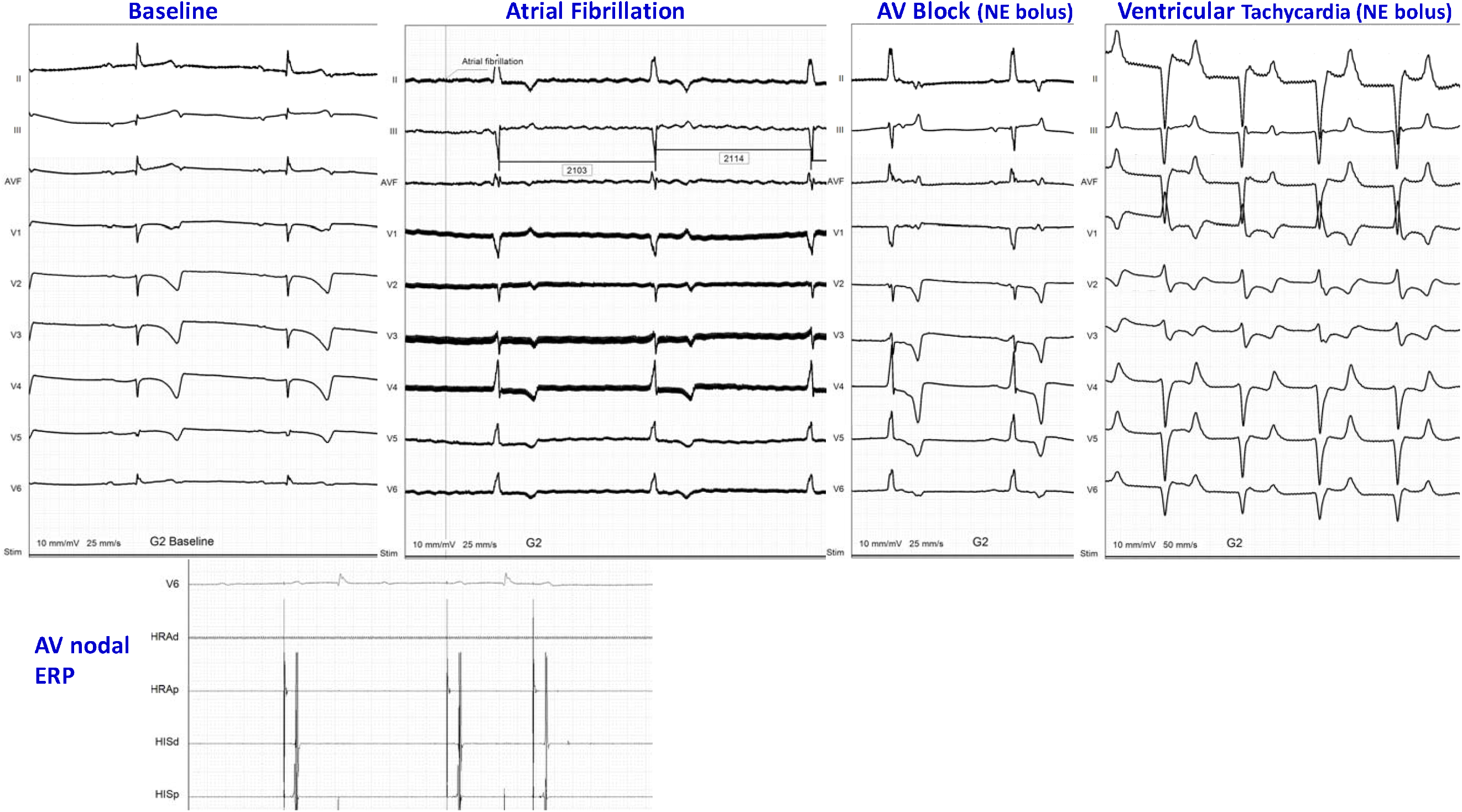
Electrophysiological study tracings from giraffe 2. The panels show the ECG at baseline, during atrial fibrillation, AV block and ventricular tachycardia. AV= atrioventricular, ERP= effective refractory period, NE= norepinephrine, HRA= high right atrium channel, His= His bundle channel

*<u>Giraffe</u> <u>3</u>*: This procedure was initially planned to be indoors but rain resulted in outdoor procedure, leading to hypothermia (temperature approximately 32-33 degrees C). Two intracardiac catheters with four platinum recording electrodes (Abbott ®) were used for recordings intracardiac signals. Underlying rhythm was sinus. RR interval of 1994 ms, PR interval of 300 ms, QRS duration of 89 ms, QT interval of 680 ms (QTc 481 ms). A His bundle signal could not be recorded with blind catheter manipulation. AV nodal ERP was 900 ms (with a basic drive train of 1600 ms). AV Wenckebach cycle length was 1500 ms. Ventricular pacing showed no VA conduction before and after NE bolus (3 mg total dose). The (LVSP) increased to 429 mm Hg from a baseline of 175 mm Hg.

An allometric scale was plotted for the PR intervals measured in this study and placed in context of the available data in the literature. The graph of PR Interval in milliseconds versus the log of body mass in kilograms for the species were used. Data was first aggregated from the available papers from the Meijler group (Meijler, 1985, Noujaim et al., 2004). The given equation for the calculated least squares regression line was graphed from Meijler (Meijler, 1985). From the Noujaim et al. (Noujaim et al., 2004), exact data for each species on PR interval and weight were plotted. To modify the data and fit it into the graph’s format of the log of body mass in kilograms versus PR interval in milliseconds, the log of all the body weights was plotted. A least squares regression line was calculated for the plotted Noujaim et al., data (Noujaim et al., 2004). The original data for giraffes 2 and 3 were plotted. As observed in the graph, both giraffes were significant outliers in their weight category in terms of PR interval, having a much longer interval compared to other species in their weight category (Figure 3).

**Figure 3:**
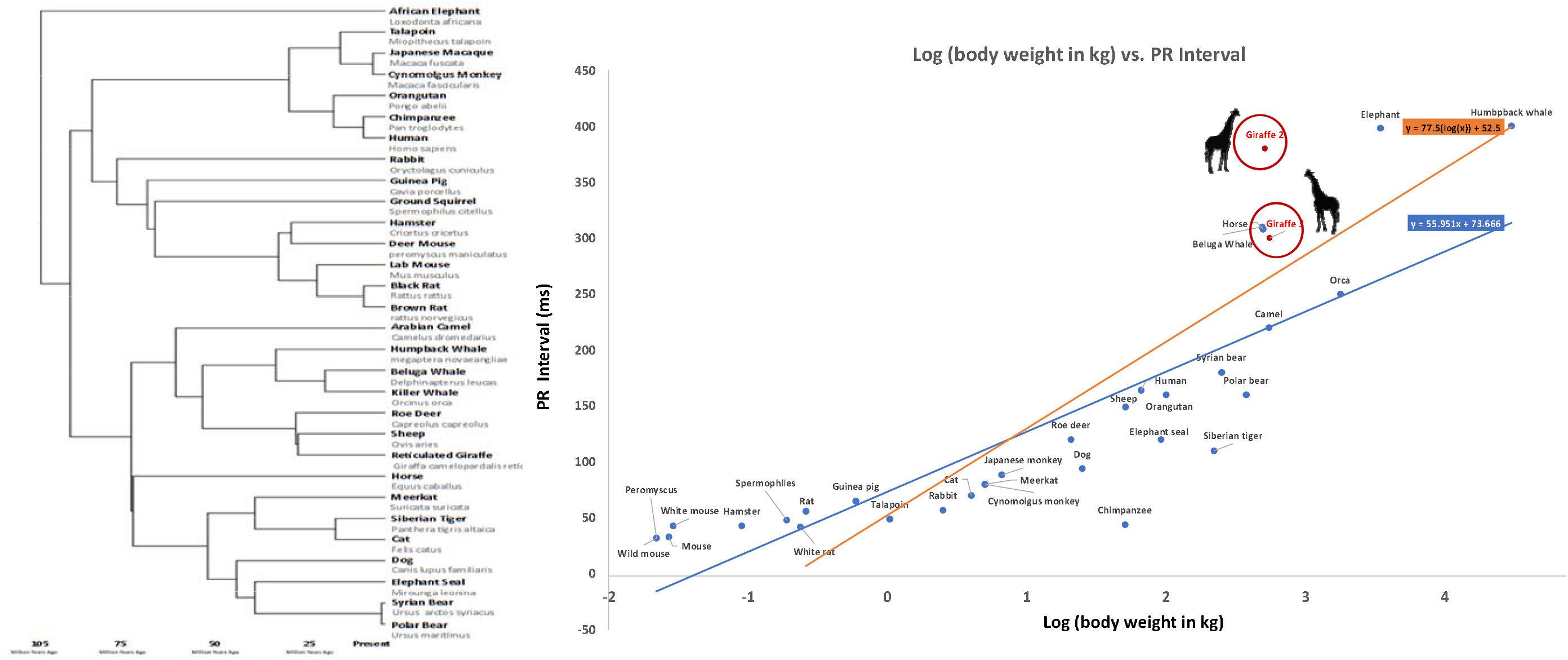
Evolutionary comparisons of PR Intervals across mammals. Left panel shows phylogenetic map of giraffes. The right panel shows the PR interval and a plot of allometric data. Giraffe PR intervals (reflecting AV nodal function) deviate from predicted allometric relationships pointing to species-specific conduction adaptations which may underlie unique characteristics of giraffe cardiovascular physiology.

### Corresponding values in a non-giraffe large ungulate (horse)

Typical corresponding values for healthy adult horse (median weight 535kg) would be a resting heart rate was 35±3 beats/min, a PR interval of 345±41 milliseconds.

## Discussion

Our ambulatory ECG data in one giraffe and invasive electrophysiological evaluation in two giraffes are the first of their kind. The ECG provides interesting insights into cardiovascular physiology and a basis for comparisons relating to specific adaptations of the cardiovascular system (Detweiler, 2010). Atrioventricular conduction has been in focus to understand how body mass affects cardiac electrical conduction (Noujaim et al., 2004), and previous measurements of PR interval in giraffes have reported an interval of 180 ms and in a wild giraffe (Goetz and Budtz-Olsen, 1955) and one in Chicago Zoo (Rossof, 1972).

The invasive approach in giraffe 2 and 3 provide insights to AV nodal effective refractory periods and the lack of ventriculoatrial (VA) conduction in both giraffes, including one where infusion of norepinephrine did not impact VA conduction. Induced atrial fibrillation was remarkable for the very slow ventricular response. The resting PR interval and slow ventricular response during atrial fibrillation could reflect significant inhibitory control of the AV nodal complex by autonomic innervation or specific electrotonic features (Meijler and Fisch, 1989) of the AV node in giraffes. Sympathetic innervation of the atria has been documented in giraffes (Nilsson et al., 1988), whereas cholinergic innervation of the giraffe heart in giraffes and the innervation of the AV nodal complex remains unknown. The lack of VA conduction is surprising and will need future studies and should be correlated to tissue features, especially innervation of the AV nodal complex.

The elevated sinus rhythm upon norepinephrine (NE) infusion resembles other mammals. The occurrence of spontaneous AV block during NE infusion was of interest and could represent a baroreflex response and a robust AV nodal inhibition by parasympathetic output. The sinus rate increased from a baseline of 1700 ms at baseline to 1118±17 ms during NE infusion that was accompanied by complete AV block with a narrow complex ventricular/junctional escape rhythm with a cycle length of 1963±53 ms. Thus, the chronotropic effect of NE infusion was accompanied by reflex AV block, which resolved when NE was discontinued. The carotid sinus is densely innervated in giraffes (Lawrence and Rewell, 1948, Nilsson et al., 1988) and heart rate is under barostatic regulation (Millard et al., 1986). A relatively narrow complex ventricular tachycardia was also seen during NE infusion that terminated spontaneously. The mechanism of this VT could not be investigated in detail, as it was self-limiting; however, it suggests an automatic rhythm from the conduction system (as the heart was normal by gross inspection at autopsy, data not shown). VA conduction was not seen during VT.

### Benefits for Giraffes and Wildlife Conservation

Anesthesia-associated mortality in giraffe may reach 10% (Bertelsen, Bush, 1987). Adverse outcomes have been associated with physiologic changes modulated by the autonomic nervous system. Current research offers insights into autonomic and conduction parameters with salience for giraffe conservation and anesthetic management.

### Benefits for Human Medicine

Species-specific adaptations in the giraffe may offer insights with salience for a heart failure with preserved ejection fraction in humans, autonomically-mediated syndromes such as takotsubo cardiomyopathy and the complications of systemic hypertension. Specifically, vagal nerve stimulation has been advanced as a therapeutic approach for heart disease.

The data from these studies may serve as a roadmap for understanding cardiac neural control and for the development of therapeutic and prevention strategies for human heart failure.

## Limitations

Some limitation of our studies includes the fact that juvenile giraffes were studied. Anesthesia and its effects are unavoidable and constitute a limitation of all invasive studies. It is important to acknowledge that data from these animals bred in captivity cannot be extrapolated in totality to giraffes in the wild. The ambient temperatures were different in giraffes that underwent invasive electrophysiological studies.

## AUTHOR CONTRIBUTIONS

BNH, TW and OAA conceived and designed the study. JH,OAA, MFB, CA MHS participated in data collection. TES performed formal analyses, curated the data and prepared the original draft of the manuscript and created the visualizations. BNH, TW, MFB, CA reviewing and editing. All authors reviewed and approved the final version of the manuscript.

## ACKNOWLEDGEMENTS

The authors thank Mr. Frank–Peter Klein, Schwarzer Cardiotek for the loaner equipment for the study and Dr. Tim Laske, Medtronic Inc for the donation of the implantable loop recorder.

